# Fourier transform infrared spectroscopy reveals high intraspecies diversity of *Malassezia pachydermatis* in dogs with atopic dermatitis

**DOI:** 10.64898/2026.04.05.716536

**Authors:** Simon Kurmann, Marco A. Coelho, Sarah Mertens, Ana Rostaher, Nina Fischer, Franco Martini, Michelle Knecht, Márcia David-Palma, Joseph Heitman, Salomé LeibundGut-Landmann, Claude Favrot, Francis Muchaamba

**Affiliations:** Section of Immunology, Vetsuisse Faculty, University of Zurich, Zurich, Switzerland; Dermatology Unit, Clinic for Small Animal Internal Medicine, Vetsuisse Faculty, University of Zurich, 8057 Zurich, Switzerland; Department of Molecular Genetics and Microbiology, Duke University School of Medicine, Durham, NC, USA; Institute of Experimental Immunology, University of Zurich, Zurich, Switzerland; Medical Research Council, Center for Medical Mycology at the University of Exeter, Department of Biosciences, Faculty of Health and Life Sciences, Exeter, UK; Institute of Food Safety and Hygiene, University of Zurich, Zurich, Switzerland

**Keywords:** *Malassezia pachydermatis*, canine atopic dermatitis, Fourier Transform Infrared spectroscopy, intraspecies diversity

## Abstract

Canine atopic dermatitis (CAD) is a chronic inflammatory skin condition sometimes associated with microbial dysbiosis, including alterations in colonization by the lipophilic yeast *Malassezia pachydermatis*. This study investigated the population diversity of *M. pachydermatis* in the ear canals of healthy and CAD-affected dogs using Fourier-transform infrared (FTIR) spectroscopy and whole genome sequencing (WGS). Among 60 dogs, *M. pachydermatis* prevalence was significantly higher in CAD cases than in healthy controls. FTIR spectroscopy revealed greater strain heterogeneity in CAD-affected dogs, often with distinct genotypes in each ear, while healthy dogs exhibited more homogeneous populations. Using a previously developed FTIR-based artificial neural network classifier, we assigned strains to three phylogroups. Strains from phylogroups I and III were significantly enriched in CAD-affected dogs, while phylogroup II was most prevalent overall and the dominant phylogroup in healthy controls. This suggests that CAD-associated inflammation may favor specific *M. pachydermatis* phylogroups and sub-clusters within phylogroups, shaping colonization dynamics. FTIR-based typing showed full concordance with WGS across 35 sequenced isolates, recapitulating relationships among phylogenetically related isolates and their similar phenotypic profiles. Overall, our findings reveal strain-level shifts in *M. pachydermatis* populations associated with CAD and establish FTIR spectroscopy as a rapid, cost-effective tool for large-scale epidemiological studies.

## 2 INTRODUCTION

*Malassezia pachydermatis* is a lipophilic yeast commonly found in the normal skin microbiota of warm-blooded animals ^1^. It has been studied most extensively in dogs. Besides its commensal lifestyle, *M. pachydermatis* is also associated with atopic dermatitis and otitis, two common manifestations in dogs affected by allergic disease ^1–3^. Canine atopic dermatitis (CAD) is one of the most common allergic diseases, affecting up to 20% of dogs, with prevalence reaching up to 50% in some breeds and increasing over time ^4,5^. Atopic dogs frequently develop otitis externa ^6^, with *M. pachydermatis* overgrowth implicated in up to 70% of cases ^7^. This condition is characterized by a waxy, moist, dark exudate, accompanied by erythema, pruritus, and lesions in the external ear canal. Besides its elevated abundance ^8,9^, *M. pachydermatis* also adopts altered activities that are thought to contribute to skin barrier disruption and inflammation, two hallmarks of atopic dermatitis ^1^. In turn, the dysregulated skin environment is thought to promote fungal overgrowth ^10–12^.

*M. pachydermatis* displays a remarkable intraspecies diversity based on multilocus sequence typing (MLST) ^13,14^, which was recently confirmed and expanded by whole genome sequencing (WGS) ^15^. This genetic diversity has been linked to functional differences between isolates ^16^. As observed in other opportunistic fungal pathogens, such genetic and phenotypic heterogeneity may translate into variations in pathogenicity and affect the fungus-host interaction ^17–21^. These observations raise the question of whether *M. pachydermatis* populations on a given skin site within a single host are monoclonal or diverse, and whether this pattern changes in the context of disease.

We recently established Fourier-Transform Infrared (FTIR) spectroscopy as a reliable method to differentiate *M. pachydermatis* strains with high resolution ^15^. FTIR spectroscopy is based on the principle that different chemical bonds within a molecule absorb infrared radiation at characteristic wavelengths, creating a unique molecular fingerprint ^22^ that reflects the biochemical composition of microorganisms, including proteins, carbohydrates, lipids, and other cellular components. Although not reaching the resolution of WGS-based methods, FTIR spectroscopy allows rapid and cost-effective strain classification. Here, we employed FTIR spectroscopy to characterize *M. pachydermatis* isolates from the external auditory canal of healthy dogs and dogs with atopic dermatitis. Specifically, we aimed to: (i) assess the diversity of *M. pachydermatis* within individual dogs, (ii) compare strain diversity between healthy dogs and those with atopic dermatitis, (iii) assess the association between *M. pachydermatis* diversity and disease severity, and (iv) classify the isolates into distinct phylogroups. Overall, this study enhances our understanding of *M. pachydermatis* diversity in dogs and provides a foundation for linking strain variation to pathogenesis and epidemiological patterns.

## 3 MATERIALS AND METHODS

### 3.1 Study participants

All dogs included in this study were sampled at the Dermatology Department of the Clinic for Small Animal Internal Medicine at the University of Zurich Animal Hospital between November 2023 and June 2024. Informed consent was obtained from all owners before enrolment, and all protocols were approved by the Cantonal Veterinary Office of Zurich (license number ZH 098_2023). The study was conducted in accordance with the guidelines of the Animal Welfare Act of Switzerland. Clinical and otoscopic examinations of all dogs were performed by the primary investigator, a licensed veterinarian.

Healthy group (n = 34): Healthy, nonpruritic dogs with no history of ear canal or skin problems were recruited during routine vaccination visits or were owned by hospital staff. Ten of these dogs were sampled while they underwent elective orthopedic procedures in the surgical department.

CAD group (n = 26): Dogs with a clinical diagnosis of CAD were included in the diseased group. The diagnosis was based on a history of atopic dermatitis and fulfilment of at least five out of eight diagnostic criteria described by Favrot et al., ^6^. To exclude other dermatological conditions, differential diagnoses such as ectoparasitic and infectious dermatoses were ruled out through standard diagnostic procedures, including skin scrapings and cytological examinations. Dogs in this group were further evaluated using the Canine Atopic Dermatitis Extent and Severity Index, version 4 (CADESI-04) ^23^, with only those scoring above 10 being included. Additionally, the severity of otitis externa was assessed using the previously published Otitis Index Score (OTIS-03), which is based on otoscopic findings ^24^. Only dogs with an OTIS-03 score greater than 2 for at least one ear were included. Detailed data for both groups are provided in **Supplementary Table S1**.

Dogs were excluded from both groups if they had received any topical ear treatment or ear cleaner within three days prior to sampling and/or systemic antifungal treatment within four weeks prior to sampling. Dogs with systemic diseases, such as autoimmune disorders, endocrine disorders, or neoplasia, were also excluded. Atopic dogs receiving concurrent medications, including lokivetmab (Cytopoint®, Zoetis Inc.), oclacitinib (Apoquel®, Zoetis Inc.), cyclosporine (Atopica®, Elanco Animal Health), or topical hydrocortisone aceponate (limited to application on the paws) were eligible for inclusion. Demographic and clinical data of the study participants are summarized in **Table 1**.

**Table 1.** Demographic and clinical data of study participants.

| Parameter<sup>a</sup> | Healthy group | CAD group |
| --- | --- | --- |
| Number of dogs | 34 [16] | 26 [17] |
| Female | 19 [8] | 12 [8] |
| Male | 15 [8] | 14 [9] |
| Male-to-Female ratio per group<sup>b</sup> | 0.79 [1] | 1.17 [1.125] |
| Median CADESI-04 score | N/A | 14 (range: 11–52) |
| Median OTIS-03 score (per ear) | N/A | 6 (range: 3–16) |
| Median age (years) | 6 (range: 1–11) | 6 (range: 2–10) |

<sup>a</sup> The table summarizes the demographic and clinical data of both healthy dogs and dogs with CAD, from which samples were collected. Numbers indicate total number of animals sampled; numbers in [] indicate number of animals from which *M. pachydermatis* could be recovered. N/A: Not applicable. <sup>b</sup> The overall male-to-female ratio of the study population is 0.94.

### 3.2 Collection of ears swabs

Ear swab samples were collected from all dogs using sterile cotton swabs (Paul Boettger GmbH & Co. KG) under standardized conditions to ensure consistency in sample collection for subsequent analyses. The swab was inserted into the external ear canal up to the junction between the vertical and horizontal canals, rotated 360 degrees, and then withdrawn to ensure sufficient sampling. Both ears of each dog were sampled separately to allow for independent analysis. Additionally, for one healthy and two CAD-affected dogs (Dogs #55, #56, and #60), two areas per ear canal were sampled: the superficial ear entrance and the deeper ear canal (between the vertical and horizontal canal), referred to as the “superficial” and “deep” swabs, respectively. Repeated sampling was conducted for one healthy dog (Dog #25; samples labelled #25 and #60) and one CAD-affected dog (Dog #30; samples labelled #30 and #50, treated with Apoquel for 3 weeks once a day), with a gap of over two months between samplings.

### 3.3 Cytological analysis for yeast cell counts

To estimate the yeast cell density in each ear, the swabs were stained with Diff-Quick for cytological analysis under ×1000 magnification using an Olympus BX53 microscope (Olympus Corporation). The average yeast count was calculated as the mean of five randomly selected microscopic fields. Not sufficient material was available for cytology for 15 healthy dogs (**Supplementary Table S1**).

### 3.4 *M. pachydermatis* isolation and culture from ear swabs

Ear swabs were plated onto modified Dixon agar (mDixon agar; 36 g malt extract, 10 g peptone, 20 g dehydrated ox bile, 10 mL Tween 40, 2 mL glycerin, 2 mL oleic acid, and 15 g agar per liter, with the pH adjusted to 6.0) for the isolation of *Malassezia* spp. ^25^. To inhibit bacterial growth, agar plates were supplemented with 1 mg/mL ampicillin. To recover additional colonies, swabs were then incubated in 2 mL liquid mDixon medium in a 10 mL sterile tube at 30°C, 180 rpm for 24 hours, after which a 10-fold concentrated culture was plated onto fresh mDixon agar supplemented with ampicillin. All plates were incubated at 30°C for 3 to 14 days and monitored for fungal growth. Then 8 individual colonies per swab were selected based on morphology (yellowish-cream coloration, smooth or slightly wrinkled surfaces, buttery consistency) ^25^ and preserved in mDixon medium supplemented with 28.6% (v/v) glycerol at −80°C. *M. pachydermatis* isolates were also grown on Sabouraud glucose agar (20 g D-glucose, 10 g peptone, and 15 g agar per liter) at 30°C for 2–5 days.

### 3.5 Other yeast strains

*M. sympodialis* strains (n=6), *M. furfur* strains (n=12), *M. globosa* strains (n=2), *M. restricta* (n=1), and *C. albicans* strains (n=2) were grown on mDixon agar and Sabouraud glucose agar at 30°C for 48 hours **(Supplementary Table S2**).

### 3.6 MALDI-TOF MS analysis

Isolates were re-streaked on mDixon agar and incubated for 48 ± 3 hours at 30°C for analysis by matrix-assisted laser desorption-ionization time-of-flight mass spectrometry (MALDI-TOF MS). Sample preparation followed to the manufacturer’s standard direct transfer protocol using α-Cyano-4-hydroxycinnamic acid as the matrix. Mass spectrometry measurements were performed with a Microflex LT MALDI-TOF mass spectrometer (Bruker Daltonics), and automated acquisition and analysis were conducted using FlexControl 3.4.206.94 and Flex Analysis 3.4.79.0 softwares (Bruker Daltonics).

### 3.7 FTIR spectroscopy analysis

Isolates were re-streaked from −80°C cryo-stocks on mDixon agar and incubated at 30°C for 2–3 days. Individual colonies were then passaged onto fresh mDixon agar and incubated for 48 ± 1 hours at 30°C. Sample preparation for FTIR spectroscopy analysis was done according to our optimized ‘water then ethanol’ protocol ^15^. Briefly, two 1 µL inoculation loops of biomass were resuspended in 70 µL deionized water in a 1.5 mL tube containing metal rods (Bruker Daltonics). After vortexing, 70 µL 70% (vol/vol) ethanol was added before 15 µL aliquots of this suspension were spotted in triplicate onto a 96-well silicon plate (Bruker Daltonics) and dried at 37°C. Infrared Test Standards IRTS 1 and IRTS 2 were also spotted in duplicates and dried under the same conditions.

Spectra acquisition was performed using the IR Biotyper^®^ system (OPUS software v8.2, Bruker Optics) with default settings. At least two independent biological replicates, each with three technical replicates, were conducted per isolate. Only spectra that passed quality control checks were used for further analysis. Spectral preprocessing included second derivative calculation, spectral region splicing, and vector normalization. Analyses focused on the carbohydrate region (1300–800 cm ¹) ^15^. For classification, Hierarchical Cluster Analysis (HCA) and Linear Discriminant Analysis (LDA) were applied. Euclidean distance and average linkage clustering methods were used to define clusters, with Cut-Off Values (COVs) determined during analysis. A dog or ear was considered to have a single cluster when most isolates grouped together, even if occasional singleton outliers were present. Results were visualized through dendrograms, distance matrices, and 2-D/3-D scatter plots. Phylogroup classification was performed using an artificial neural network (ANN)-based FTIR spectroscopy classifier previously developed ^15^.

### 3.8 DNA extraction, whole-genome sequencing, and bioinformatic analysis

Genomic DNA was extracted using the Quick-DNA Fungal/Bacterial Miniprep Kit (Zymo Research), following the manufacturer’s instructions, using 50 µL elution buffer, and quantified with a NanoDrop spectrophotometer and a Quantus Fluorometer (Promega). DNA integrity was assessed with a BioAnalyzer (Agilent Technologies), and samples were stored at −80°C until further processing. Libraries were prepared with TruSeq® Nano DNA Library Prep (Illumina) and sequenced at the Functional Genomics Centre Zurich on an Illumina NovaSeq X Plus platform to generate 2×151 bp paired-end reads.

De novo assemblies were generated with SPAdes v4.2.0, retaining only scaffolds >500 bp to minimize inclusion of short, likely spurious contigs. Assembly statistics (genome size, scaffold count, N50/L50, N90/L90, GC content, and coverage) were computed using a custom Python script and are summarized in **Supplementary Table S3**. Ploidy was inferred from total assembly length relative to previously characterized haploid genomes of *M. pachydermatis* ^26–28^.

To assess lineage diversity, FastANI v1.34 was used to calculate an all-vs-all pairwise average nucleotide identity (ANI) matrix across all isolates and representative genomes from the three known *M. pachydermatis* phylogroups (PG I: KCTC27587 and CBS1879; PG II: MA165 and MA718; PG III: MA9 and MA1174). Custom Python scripts converted ANI values into distance scores (100 – ANI; **Supplementary Table S4**), applied UPGMA (average-linkage) clustering, and generated Newick-format dendrograms. Phylogroup assignment was based on clustering with the corresponding reference genomes in the ANI dendrogram. Resulting dendrograms were visualized with iTOL v7. WGS-based phylogroups were then compared to those assigned by the FTIR spectroscopy ANN classifier to evaluate concordance.

### 3.9 Phospholipase activity analysis

Phospholipase (PLA2) activity was evaluated by means of the semi-quantitative egg-yolk method described previously ^29,30^. Briefly, 5 μL of a fresh *M. pachydermatis* culture containing 10^7^ cells/mL was spotted on Price’s egg yolk agar (40 g glucose, 10 g peptone, 58.44 g NaCl, 0.277 g CaCl_2_ and 17 g agar per liter with 80 mL egg yolk emulsion (Sigma) added after autoclaving). Plates were incubated for 28 days at 37°C in a sealed plastic box to maintain humidity. The formation of a precipitation zone around the colonies was indicative of PLA2 activity. The PLA2 activity index (Pz) was calculated as the ratio of the total diameter (colony + precipitation zone) to colony diameter ^29^. Samples with a Pz value equal to 1.0 were considered negative for PLA2 activity.

### 3.10 Statistical analysis

FTIR spectra were analyzed using HCA and LDA, with clusters defined by Euclidean distance and average linkage clustering. Comparisons between clusters derived from FTIR spectroscopy and those generated by WGS, based on ANI, were performed using Simpson’s Index of Diversity (SID), the Adjusted Rand Index (ARI), and the Adjusted Wallace Coefficient (AWC). These metrics, which evaluate diversity (SID), overall agreement (ARI), and directional congruence (AWC) between clustering methods, were calculated with 95% confidence intervals using an online tool (http://www.comparingpartitions.info/). ARI and AWC values range from 0 to 1, where 0 indicates agreement expected by chance, and 1 indicates perfect correlation between methods ^31–33^. WGS was regarded as the gold standard for these comparative analyses.

Statistical analysis of non-FTIR spectroscopy data was conducted using GraphPad Prism (Version 9.2.0, GraphPad Software). Fisher’s exact test and Mann-Whitney U test were used to assess associations between categorical variables. PLA2 activity data (Pz values) were assessed for normality using the Shapiro–Wilk and Anderson–Darling tests, and homogeneity of variances was evaluated with Levene’s test. Depending on data distribution, analyses used one-way ANOVA with Tukey’s post-hoc tests or Kruskal-Wallis test followed by Dunn’s post-hoc t-tests. For comparisons between two groups (e.g., mean PLA2 activity of isolates from the left versus right ear), an unpaired t test with Welch’s correction or the Mann–Whitney U test was applied, as appropriate. P < 0.05 was considered statistically significant.

### 3.11 Data availability statement

All raw data linked to this study will be made publicly available at zenodo.org upon acceptance of the manuscript (doi will be provided). Raw sequencing reads, and genome assemblies were deposited in DDBJ/EMBL/GenBank under BioProject PRJNA1402406.

## 4 RESULTS

### 4.1 Ear colonization by *Malassezia pachydermatis* in healthy and CAD dogs

To establish a collection of canine *M. pachydermatis* isolates, we sampled 34 healthy dogs and 26 dogs diagnosed with atopic dermatitis (**Table 1**). Both ears of each dog were sampled individually, resulting in 68 swabs from healthy dogs and 52 swabs from CAD-affected dogs. Eight representative colonies from each positive swab were selected for further analysis. Initial identification was based on colony morphology and the ability of isolates to grow on Sabouraud glucose agar, whereas non-*pachydermatis* species of *Malassezia* failed to grow under these conditions. The species identity of representative isolates was further assessed by MALDI-TOF MS, and six swabs were excluded as the recovered isolates were not confirmed as *M. pachydermatis* (**Supplementary Table S1**).

In the healthy control group, *M. pachydermatis* was recovered from 27/68 swabs (39.71%), originating from 16 dogs (dog-level prevalence: 16/34 dogs, 47.06%) (**Figure 1**). Among these 16 dogs, 11 tested positive in both ears, whereas the remaining five tested positive in only one ear. In contrast, the isolation rate of *M. pachydermatis* in the CAD group was significantly higher (34/52 swabs, 65.38%; P = 0.0061, Fisher’s exact test), with 17/26 dogs (65.38%) testing positive (**Figure 1**). All 17 CAD dogs exhibited bilateral ear involvement. In total, the collection comprised 200 *M. pachydermatis* isolates from healthy dogs and 259 isolates from CAD dogs. Additionally, 32 isolates were obtained from repeated sampling (n = 2 dogs; one healthy and one CAD-affected) and 48 from superficial ear-site sampling (n = 3 dogs; one healthy and two CAD-affected). In total, this yielded 539 isolates for FTIR-based characterization.

**Figure 1.**
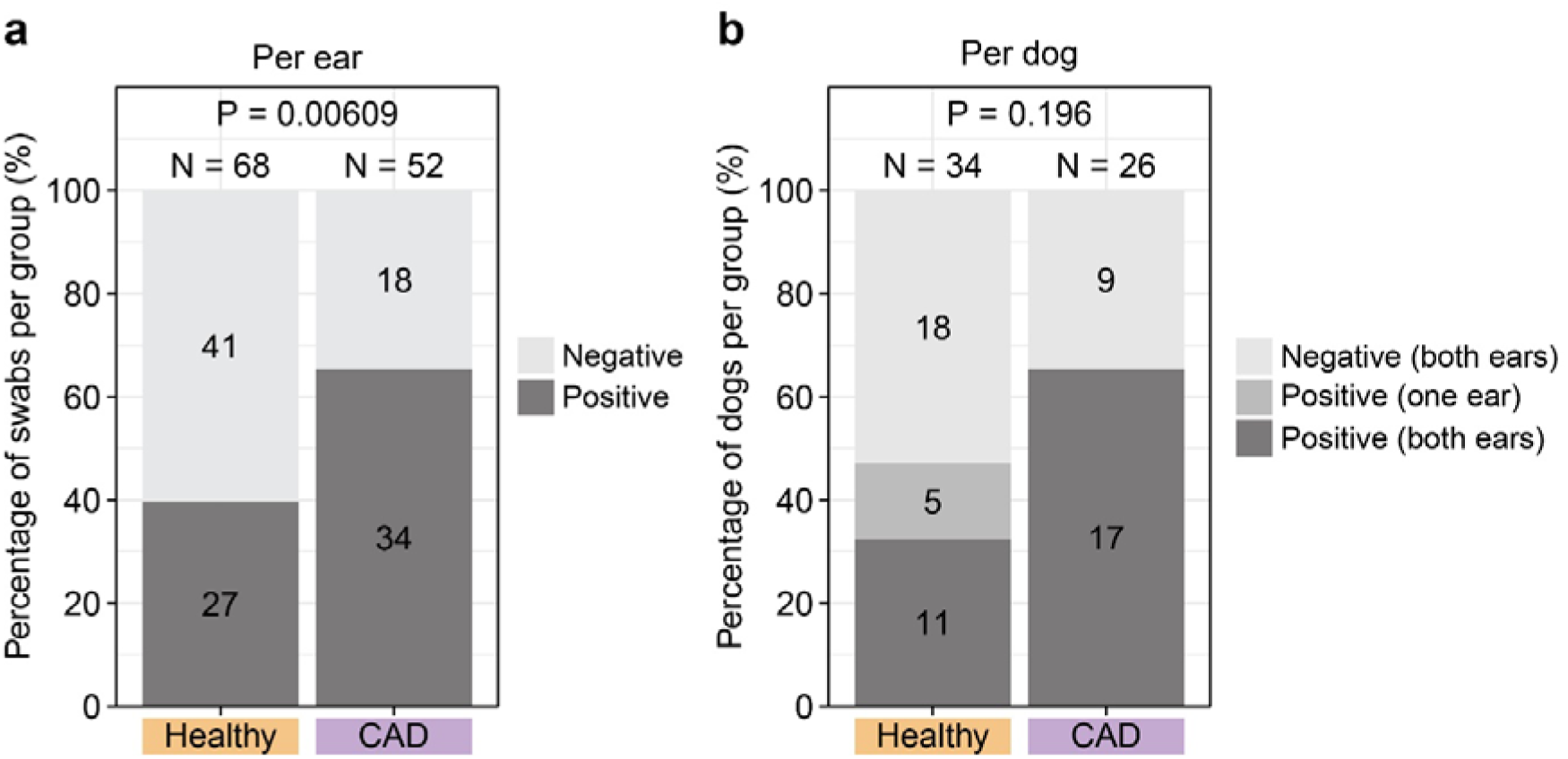
Comparison of *M. pachydermatis* isolation rates between healthy and CAD-affected dogs. **(a)** Percentage of ear swabs testing positive (dark grey) or negative (light grey), relative to the total number of swabs in each group. **(b)**. Percentage of dogs testing positive in both ears (dark grey), positive in one ear only (middle grey), or negative in both ears (light grey), relative to the total number of dogs in each group. Numbers within bars indicate the absolute numbers of ears swabs **(a)** or dogs **(b)** used to calculate percentages. Statistical analyses were performed on the raw data (absolute counts) with Fisher’s exact test.

### 4.2 FTIR spectra of *M. pachydermatis* isolates from healthy and CAD dogs

All 539 *M. pachydermatis* isolates were examined by FTIR spectroscopy to determine the similarity between isolates. FTIR spectroscopy discriminates *M. pachydermatis* from other *Malassezia* spp. such as *M. furfur*, *M. globosa*, *M. restricta* and *M. sympodialis* ^15^. All *M. pachydermatis* isolates clustered closely together and were clearly distinct from any of the other *Malassezia* species included in the experiment and from *C. albicans*, which was included as a control (**Figure 2, Supplementary Figure S1**).

**Figure 2.**
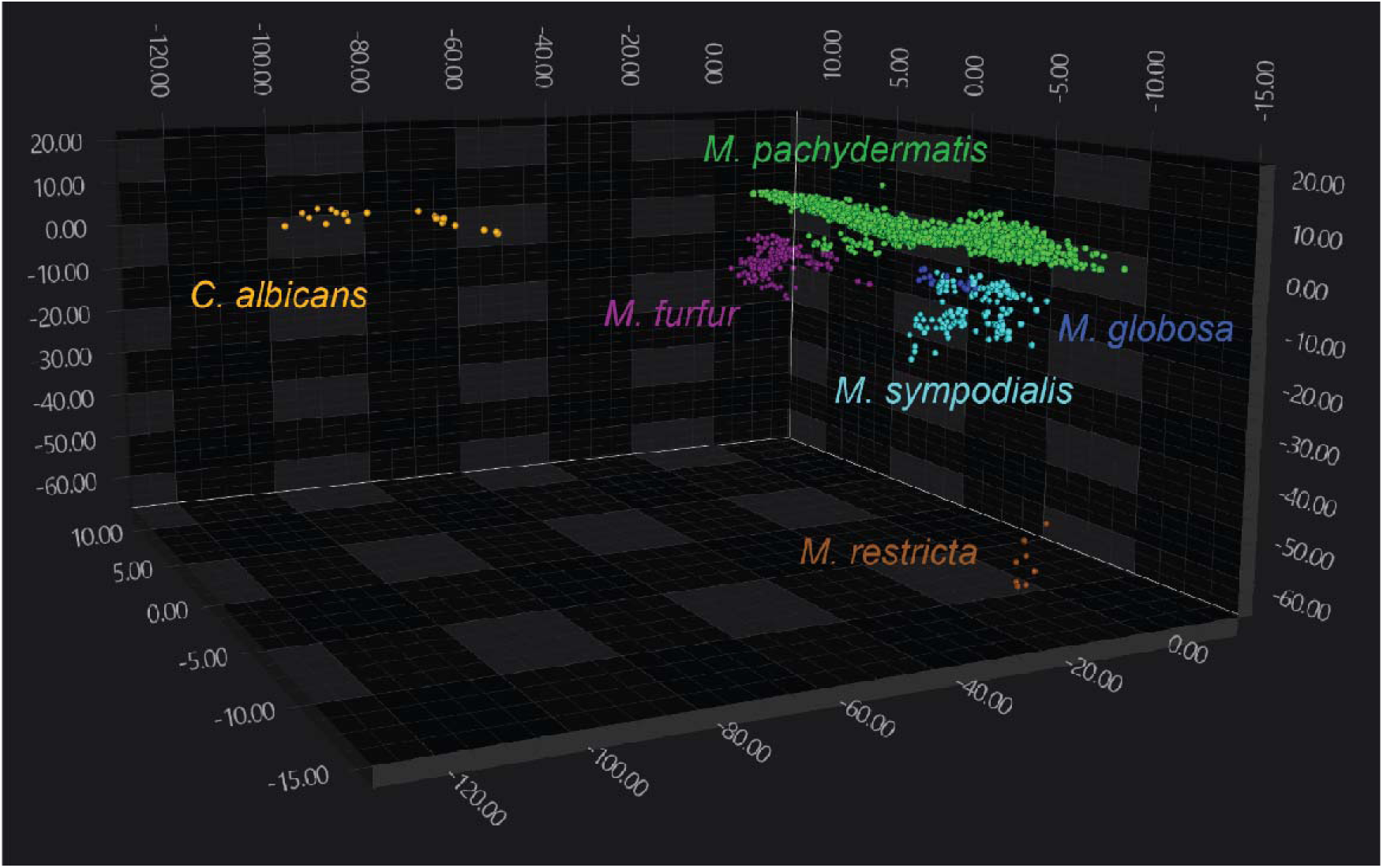
Separation of *M. pachydermatis* from other *Malassezia* species in the FTIR spectral space. 3D scatter plot showing the distribution of *M. pachydermatis* isolates from study dog ears (n = 539) and reference strains of *Malassezia furfur* (n = 12), *M. globosa* (n = 2), *M. sympodialis* (n = 6), *M. restricta* (n = 1) and *C. albicans* (n = 2) within the FTIR spectral space. The plot displays the first three LD axes. Each isolate is represented by at least six spectra, with each symbol (•) corresponding to a technical replicate from at least two independent biological repeats, i.e., each isolate is represented by at least 6 symbols. Dimensionality reduction was performed using LDA.

When further analyzing the FTIR spectra of the *M. pachydermatis* isolates, we noticed a distribution into three main clusters (**Figure 3**). Each cluster contained isolates from both healthy dogs and dogs with CAD. However, within each cluster, the two groups of isolates were not evenly distributed. Isolates from CAD-affected dogs showed a tendency to cluster separately from those of healthy dogs especially in cluster 2 (**Figure 3**). These findings suggest that, while overlap between isolate groups exists, CAD isolates may exhibit distinct traits in association with their role in disease.

**Figure 3.**
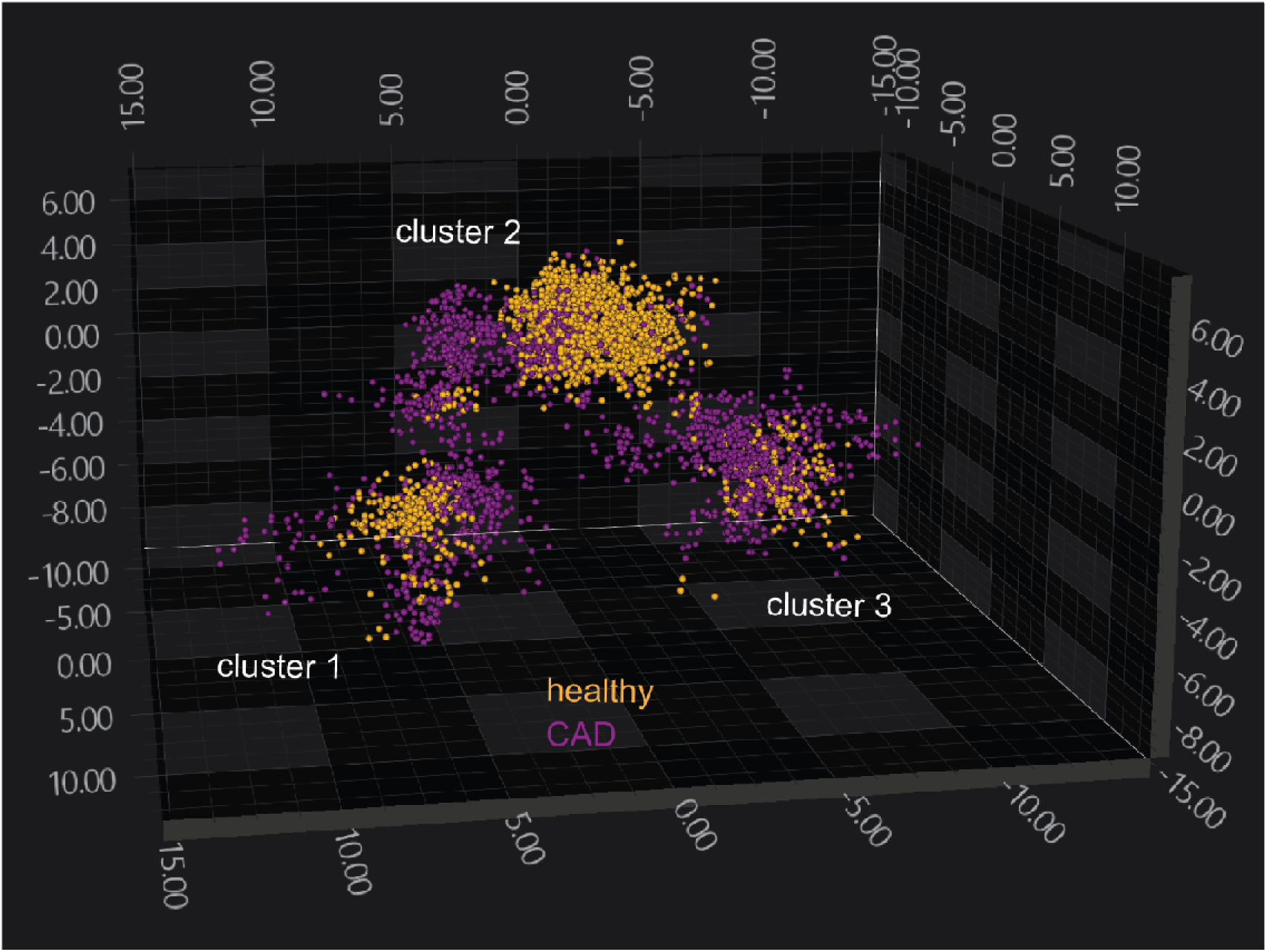
*M. pachydermatis* clustering based on the health status of the dog from which they were obtained. 3D scatter plot showing the distribution of *M. pachydermatis* isolates from ears of healthy (orange) and CAD-affected dogs (purple) within the FTIR spectral space. The plot displays the first three LD axes. Each isolate is represented by at least six spectra, with each symbol (•) corresponding to a technical replicate from at least two independent biological repeats, i.e., each isolate is represented by 6 symbols. Dimensionality reduction was performed using LDA

### 4.3 *M. pachydermatis* isolate distribution in healthy and diseased dogs

Next, we evaluated the distribution of isolates in individual ears and dogs across the entire study population. To do this, we analyzed FTIR spectroscopy clusters determined from distance matrices using automatically calculated cut-off values. When examining the clustering patterns per ear, we observed that in most ears (65.57%, 40/61), the entire population clustered within a single group, suggesting that all isolates were closely related and consistent with colonization by one dominant strain, whereas in the remainder of ears (34.43%, 21/61) isolates were distributed across two clusters (**Figure 4**). Focusing on healthy dogs, isolates from 74.1% (20/27) of ears constituted a single cluster per ear, while 25.9% (7/27) were split into two clusters (**Figure 4A, left**). In the CAD group however, the pattern was shifted, with a comparatively smaller proportion of ears containing a single cluster (58.8%, 20/34) and a larger proportion containing two clusters (41.2%, 14/34) (**Figure 4A, left**). Notably, no ear in either group exhibited more than two clusters, although some ears with two clusters also contained singleton outliers.

**Figure 4.**
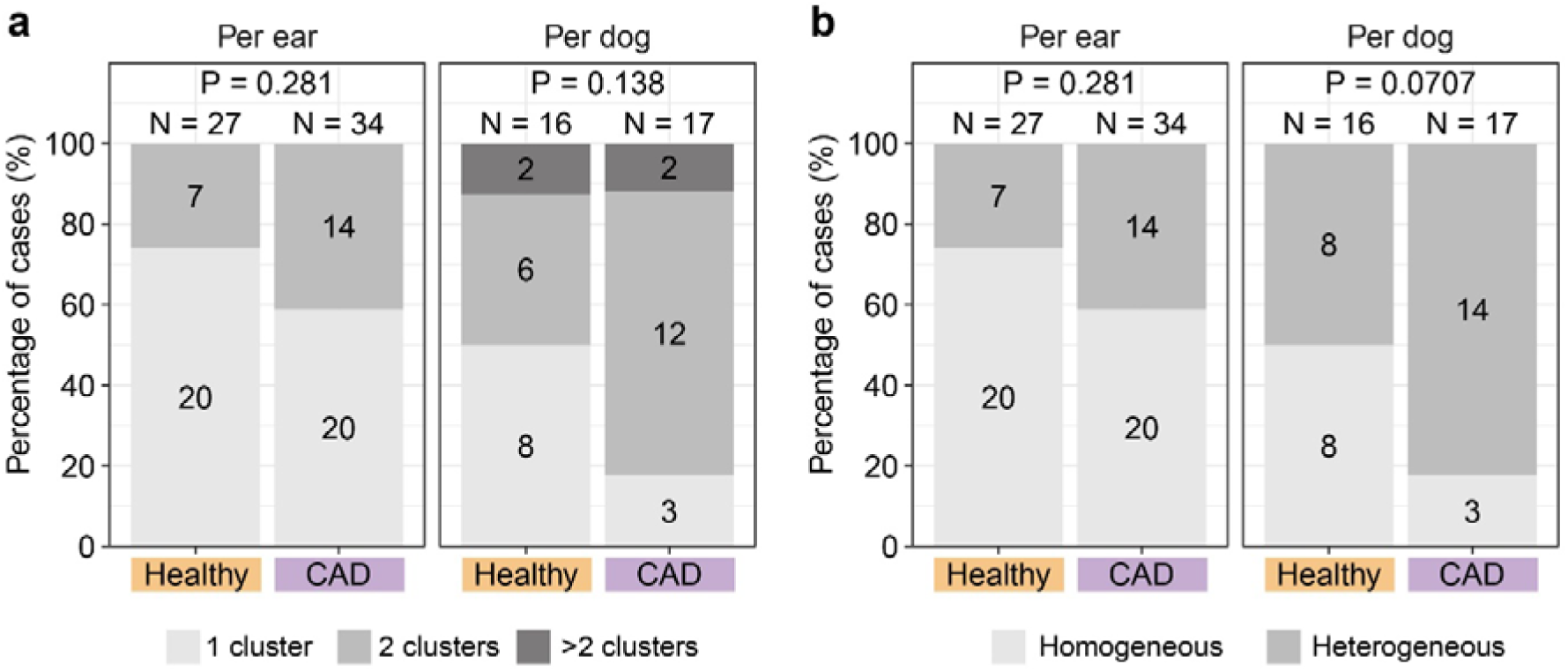
Clustering patterns and colonization complexity of *M. pachydermatis* in healthy and CAD dogs based on FTIR profiles. Bars represent the percentage of cases in healthy and CAD-affected dogs relative to the total number in each group. **(a)** Proportion of ears (left) and dogs (right) with one (light grey), two (middle grey), or more than two (dark grey) clusters. **(b)** Distribution of colonization types per ear (left) and dog (right), classified as homogeneous (1 cluster) or heterogeneous (≥2 clusters). Statistical analysis were performed on the raw data (absolute counts) with Fisher’s exact test. Because no ear samples showed more than two clusters, the comparisons shown for ears in panels A and B yield the same p-value.

A comparison of isolate diversity between the right and left ears of individual dogs revealed four recurring patterns. In some dogs, isolates from both the right and left ears clustered closely, suggesting colonization by a single fungal strain (e.g., Dog #2; scenario A, **Figure 5a**). However, in most dogs, distinct diversity patterns were observed between the two ears. For instance, in Dog #41, each ear harbored a homogeneous population, yet these populations were distinct between the right and left ears (scenario B, **Figure 5b**). Similarly, Dog #30 exhibited a distinct population in each ear, although the population from the left ear was subdivided into two separate clusters (scenario C, **Figure 5c**). In Dogs #16 and #46, we observed partial overlap in the populations from the right and left ears (scenario D). While one ear harbored a homogeneous population, the other ear displayed a heterogeneous distribution of the isolates, with some clustering together with isolates from the opposite ear (**Figure 5d-e**). The different clustering pattern were also evident in dendrograms generated from the average FTIR spectrum of each isolate, which clearly reflect the ear-specific differences in fungal diversity (**Figures 5f-j**).

**Figure 5.**
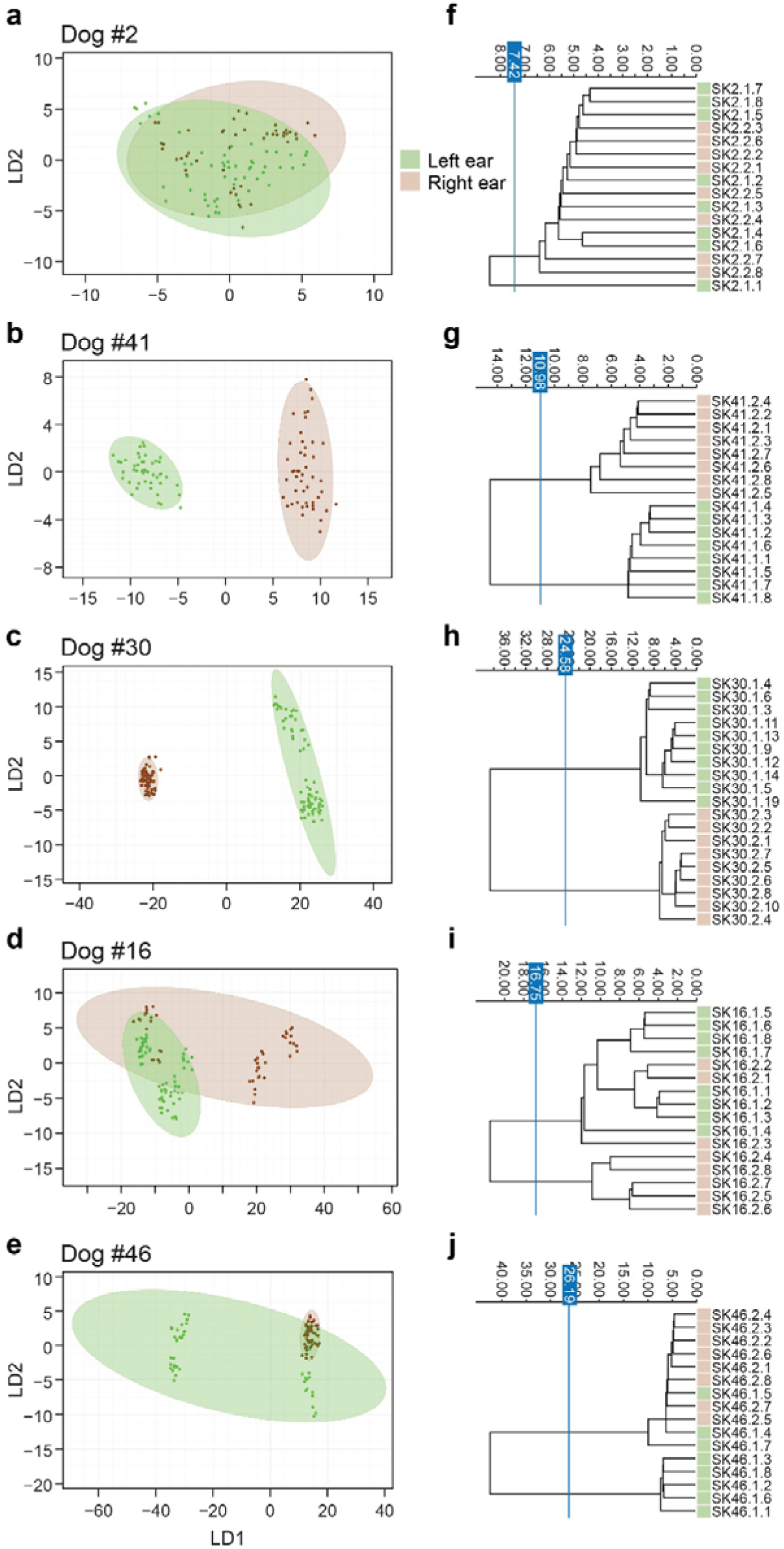
Within-dog diversity of *M. pachydermatis* isolates. Scatter plots (**a–e**) and corresponding dendrograms (**f–j**) showing the distribution of *M. pachydermatis* isolates from the left (lime) and right (brown) ears of the indicated dogs within the FTIR spectral space. For each dog, 16 isolates were analyzed (8 per ear). Each isolate is represented by at least six spectra, obtained from three technical replicates across a minimum of two independent biological replicates. Data were processed using LDA, and the scatter plots (**a–e**) show the first two LD axes. The dendrograms (**f–j**) display the average spectrum for each isolate. The vertical blue line denotes the automatically determined clustering cut-off.

Overall, of the 16 healthy dogs, isolates from 8/16 dogs (50%) clustered homogenously according to scenario A, either with or without singleton outliers. In another 6 healthy dogs (37.5%), isolates grouped into two clusters, and in the remaining 2/16 (12.5%) dogs, isolates grouped in more than two clusters (**Figure 4a, right**). In contrast, the majority of CAD-affected dogs (14/17, 82.4%) exhibited two or more clusters. Together, these results highlight differences in colonization complexity between healthy and CAD-affected dogs, with more heterogeneous colonization patterns predominating in the CAD group (**Figure 4b**), although these differences did not reach statistical significance, likely due to the limited sample size. Among the dogs with bilateral ear colonization, most (19/28, 67.9%) exhibited heterogeneous colonization, characterized by distinct clusters in opposite ears. Differences between homogenous vs. heterogenous isolate clusters in CAD dogs did not correlate with disease severity based on CADESI-04, OTIS-03, and cytology scores (**Table 2**). Of note, OTIS-3 and cytology scores are not exact and quantitative measures.

**Table 2.** Colonization complexity as a function of disease severity.

| Metric | Homogeneous population<br>(1 cluster)<sup>a</sup> | Heterogeneous population<br>(≥2 clusters)<sup>a</sup> | p-value<sup>b</sup> |
| --- | --- | --- | --- |
| CADESI-04 | 13 ± 2.83 | 17.38 ± 8.52 | 0.3366 |
| OTIS-03 | 5.95 ± 3.02 | 6.8 ± 4.5 | 0.5341 |
| Cytology | 15.38 ± 10.21 | 12.1 ± 12.14 | 0.426 |

<sup>a</sup> Mean ± standard deviation. <sup>b</sup> Based on Fisher's exact test

### 4.4 Longitudinal and site-specific comparison of *M. pachydermatis* isolates

To assess the impact of the sampling site on the clonal distribution of *M. pachydermatis* isolates, we sampled three dogs at a superficial ear site, in addition to the routinely sampled deep ear site. A direct comparison of the isolates from the two sampling sites within the same ear revealed that they often differed (**Supplementary Figures S2**). For example, the isolates from the superficial swabs of Dogs #55 and #60 (right ears) formed distinct subclusters from those obtained from the corresponding deep ear swabs (**Supplementary Figures S2b and S2h**). However, in some instances (e.g., Dogs #56), isolates from superficial and deep sites clustered together, suggesting partial overlap in microbial composition between the two sites (**Supplementary Figure S2d-f**). No consistent pattern indicating greater isolate diversity at either the superficial or deep site was observed within the same ear. Overall, deep ear swabs provided a more comprehensive representation of microbial diversity present at the lesional site, suggesting that it is the more informative sampling site.

We also had the opportunity to sample one healthy and one diseased dog longitudinally, with an interval of more than two months between the two samplings. In both cases, the *M. pachydermatis* isolate population changed over time (**Figure 6, Supplementary Figure S3**). In the healthy Dog #25, entirely new homogeneous clusters emerged over the two-month interval, and the original cluster was no longer detected (**Figure 6a, Supplementary Figure S3a-b**).

**Figure 6.**
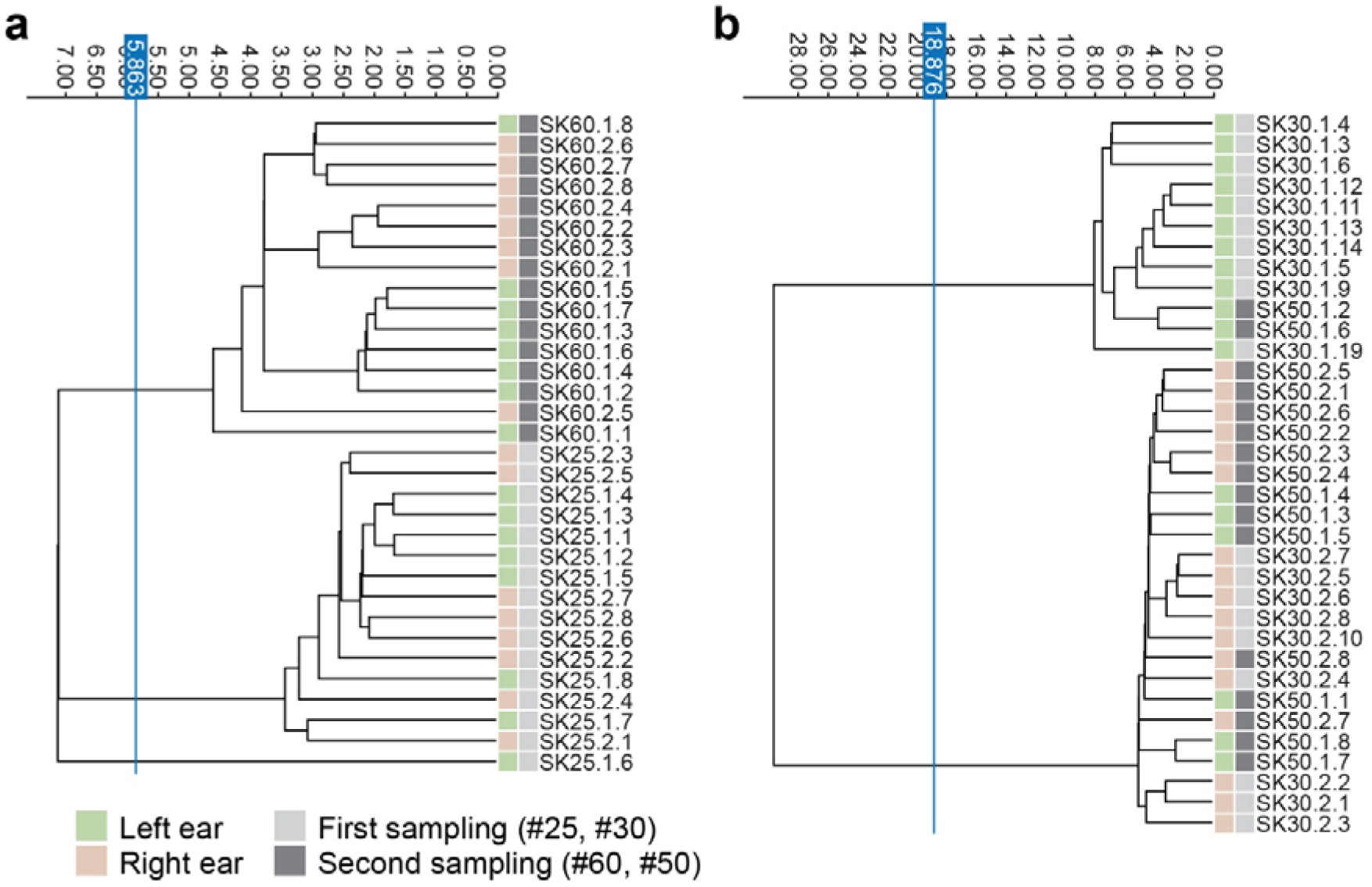
Shifts in *M. pachydermatis* populations over time. Dendrograms generated from clustering FTIR spectra of *M. pachydermatis* isolates recovered from a healthy dog (**a**; Dog #25) and a CAD-affected dog (**b**; Dog #30), each sampled twice over an interval of more than two months. Clustering was based on the average spectra from at least two independent biological replicates, each with three technical replicates. The vertical blue line denotes the automatically calculated clustering cut-off.

By contrast, in CAD-affected Dog #30, the same isolate clusters persisted over time, although their relative representation changed (**Figure 6b, Supplementary Figure S3c-d**). Two clusters were maintained in the left ear, while the dominating strain in the right ear appeared to expand into the left ear over time. Meanwhile, the right ear retained comparable isolate diversity, indicating stable, homogeneous colonization. Overall, these changes resulted in a modest shift from a heterogeneous to a more homogeneous population over time. Clinical scores reflected minor changes in the health status of the animal, with the CADESI-04 score decreasing slightly from 29 to 27, while the OTIS-03 score remained unchanged at 3 between the first and the second sampling time points.

### 4.5 FTIR spectroscopy-based phylogroup classification of *M. pachydermatis* isolates

Next, we used an artificial neural network (ANN)-based classifier developed in our previous study ^15^ to assign the *M. pachydermatis* isolates from this study to distinct phylogroups. The three major clusters observed in **Figure 3** corresponded to the three previously defined phylogroups, with cluster 1 mapping to phylogroup I, cluster 2 to phylogroup II, and cluster 3 to phylogroup III. The majority of the isolates were assigned to phylogroup II (215/491; 43.8%), followed by phylogroup I (106/491; 21.5%) and phylogroup III (85/491; 17.3%) (**Figure 7** and **Table 3**). The remaining 85 isolates could not be confidently assigned to a single phylogroup, including 69 ambiguously classified between phylogroups II and III and 16 from Dog #38 between phylogroups I and II. Phylogroup II comprised the largest fraction of isolates from healthy dogs (144/215; 67%; 18 ears). In contrast, isolates from CAD-affected dogs were more evenly distributed across all three main phylogroups. Accordingly, phylogroups I (82/106; 77.4%) and III (69/85; 81.2%) were predominantly composed of isolates from CAD-affected dogs (**Figure 7** and **Supplementary Table S5**).

**Figure 7.**
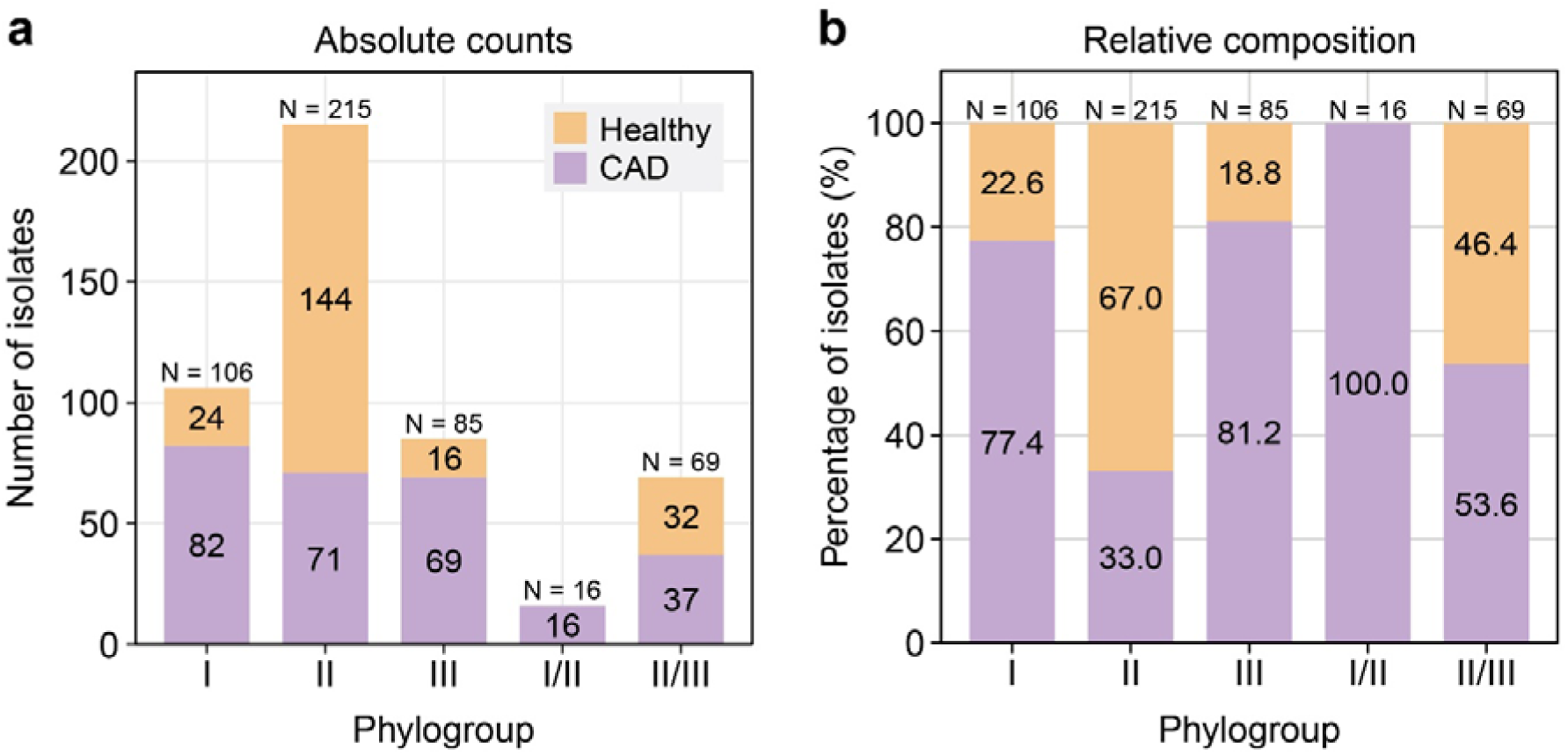
Distribution of *M. pachydermatis* isolates across phylogroups. **(a)** Absolute number of isolates assigned to each phylogroup, shown as stacked bars according to whether isolates originated from CAD-affected (purple) or healthy (orange) dogs. **(b)** Relative composition of each phylogroup, expressed as the percentage of isolates from CAD-affected and healthy dogs. Numbers within bars indicate absolute counts in panel **(a)** and percentages in panel **(b)**, whereas values above bars indicate the total number of isolates in each phylogroup.

A detailed analysis of isolates from individual dogs revealed several instances of mixed or shifting phylogroup profiles (**Supplementary Table S6 and S7**). For instance, Dog #30 was colonized with isolates belonging to different phylogroups in each ear, with phylogroup I detected in the left ear and phylogroup II in the right ear. Resampling after two months (Sample #50) revealed a mixed phylogroup profile in the left ear of Dog #30, with two isolates (SK50.1.2 and SK50.1.6) belonging to phylogroup I and the remaining six isolates assigned to phylogroup II, suggesting a potential replacement of the initial phylogroup over time. Dog #54’s left ear also exhibited a mixed pattern, with most isolates classified as phylogroup II/III, while three isolates (SK54.1.2, SK54.1.7, and SK54.1.8) belonged to phylogroup I. Similarly, Dog #46 displayed a mixed phylogroup composition in the left ear, with five isolates classified as phylogroup III and three as phylogroup I (SK46.1.4, SK46.1.5, and SK46.1.7). For Dog #60, the isolates from the right ear superficial swab were part of phylogroup III, while those from the corresponding deep swab belonged to phylogroup II. In contrast, all isolates from both the deep and superficial swabs of the left ear were classified as phylogroup II. Overall, Dogs #30, #43, #46, #50, and #54 harbored different phylogroups in opposite ears, highlighting the potential for ear-specific microbial dynamics within individual hosts (**Supplementary Table S6 and S7**).

### 4.6 Validation of FTIR spectroscopy-based observations

To validate the clustering predicted by FTIR spectroscopy, a subset of 35 isolates from four dogs was selected for WGS. Isolates were chosen to represent all three phylogroups and to capture the range of within-dog diversity scenarios inferred from FTIR analyses (**Figure 5**), allowing us to directly assess whether FTIR clustering accurately reflects underlying genetic diversity within individual dogs. High-quality genome assemblies were obtained for all sequenced isolates, with scaffold N50 values ranging from 687,718 to 1,431,285 bp and total assembly sizes consistent with the expected haploid genome size of the species (**Supplementary Table S3**) ^26–28^. FastANI comparisons confirmed that the FTIR spectroscopy-predicted phylogroup assignments per isolate were fully concordant with the WGS-based classifications (**Figure 9** and **Supplementary Figure S4**), with both ARI and AWC equal to 1.00. The within-dog and within-ear clusters identified by both WGS and FTIR (**Figure 5** and **Supplementary Table S8**), were likewise fully concordant (ARI = 1.00; AWC = 1.00), indicating that FTIR can reliably detect within-dog and within-ear diversity. When combining WGS and FTIR data from isolates across all four dogs, agreement at the cluster or subcluster level (**Supplementary Figure S4** and **Supplementary Table S8**) remained good, with an ARI of 0.630. Directional comparison using the AWC showed that FTIR clusters strongly predicted WGS clusters [AWC _FTIR → WGS_ = 0.877 (0.787-0.968)], whereas WGS clusters predicted FTIR clusters with lower agreement [AWC _WGS → FTIR_ = 0.491 (0.366-0.617]. This asymmetry likely reflects both the higher discriminatory resolution of WGS and the fact that FTIR captures phenotypic rather than strictly genomic variations, resulting in some WGS-distinct isolates being grouped within the same FTIR cluster.

The WGS analysis further supported the FTIR spectroscopy-predicted patterns of homogeneous and heterogeneous colonization at both the ear and dog levels. In Dogs #16 and #46, WGS corroborated the observed overlap in the populations from the right and left ears, with shared genetic lineages detected across ears (scenario D; **Figure 5 and 8, Supplementary Table S8**). Additionally, Dog #16 was colonized by two closely related yet genetically distinct lineages, one comprising four isolates and the other seven isolates, both belonging to Phylogroup II. Within-lineage divergence was minimal (≤0.00333%), while between-lineage divergence was significantly higher (mean 0.08248%), allowing clear separation at the within-host scale (**Supplementary Table S4**). A similar pattern was observed in Dog #41, which was colonized by two closely related lineage, also both assigned to phylogroup II. Notably, isolates SK41.2.1, SK41.2.2, and SK41.2.8 from Dog #41 clustered closely with isolates SK16.1.1, SK16.1.2, SK16.2.1 and SK16.2.2 from Dog #16, suggesting potential strain sharing or recent common ancestry across hosts **(Figure 8, Supplementary Table S8**). In contrast, Dog #46 was colonized by two lineages belonging to different phylogroups (I and III), with the left ear harboring both lineages, including four isolates assigned to phylogroup III and two to phylogroup I. WGS analysis also confirmed that Dog #30 harbored distinct phylogroups in opposite ears, with phylogroup I in the left ear and phylogroup II in the right ear, consistent with FTIR spectroscopy inferences.

**Figure 8.**
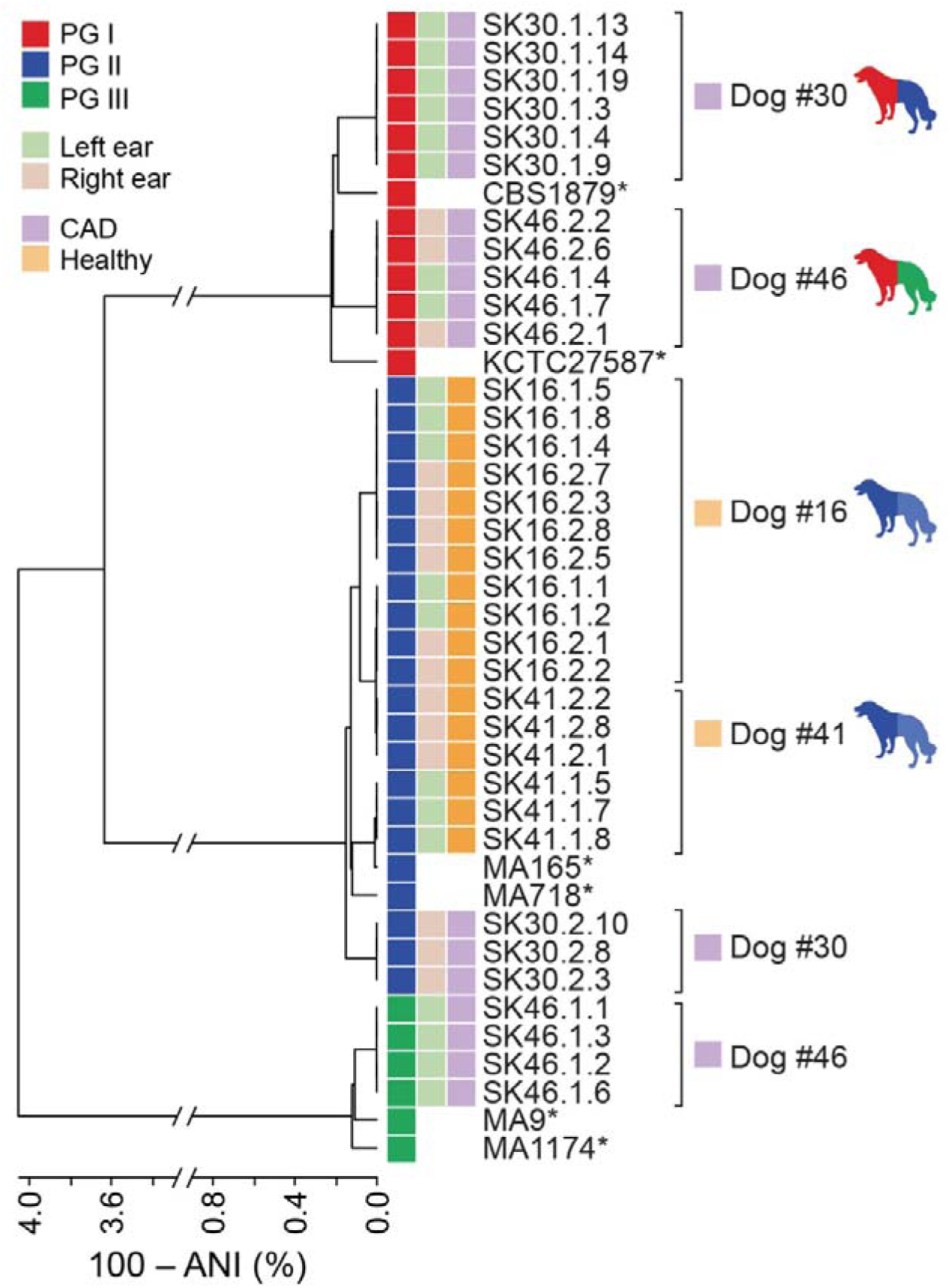
Hierarchical clustering of *M. pachydermatis* strains by WGS. Hierarchical clustering of 35 *M. pachydermatis* isolates. Two reference strains were included for each phylogroup (*). Pairwise average nucleotide identity (ANI) values were calculated using FastANI, and the resulting distance matrix was used to generate a hierarchical clustering dendrogram. Isolate identifiers follow the format SK.DogID.EarSide.IsolateNumber; the middle digit indicates the ear side, where 1 = left ear and 2 = right ear (e.g., SK16.1.1 = Dog 16, left ear, isolate 1; SK16.2.1 = Dog 16, right ear, isolate 1).

**Figure 9.**
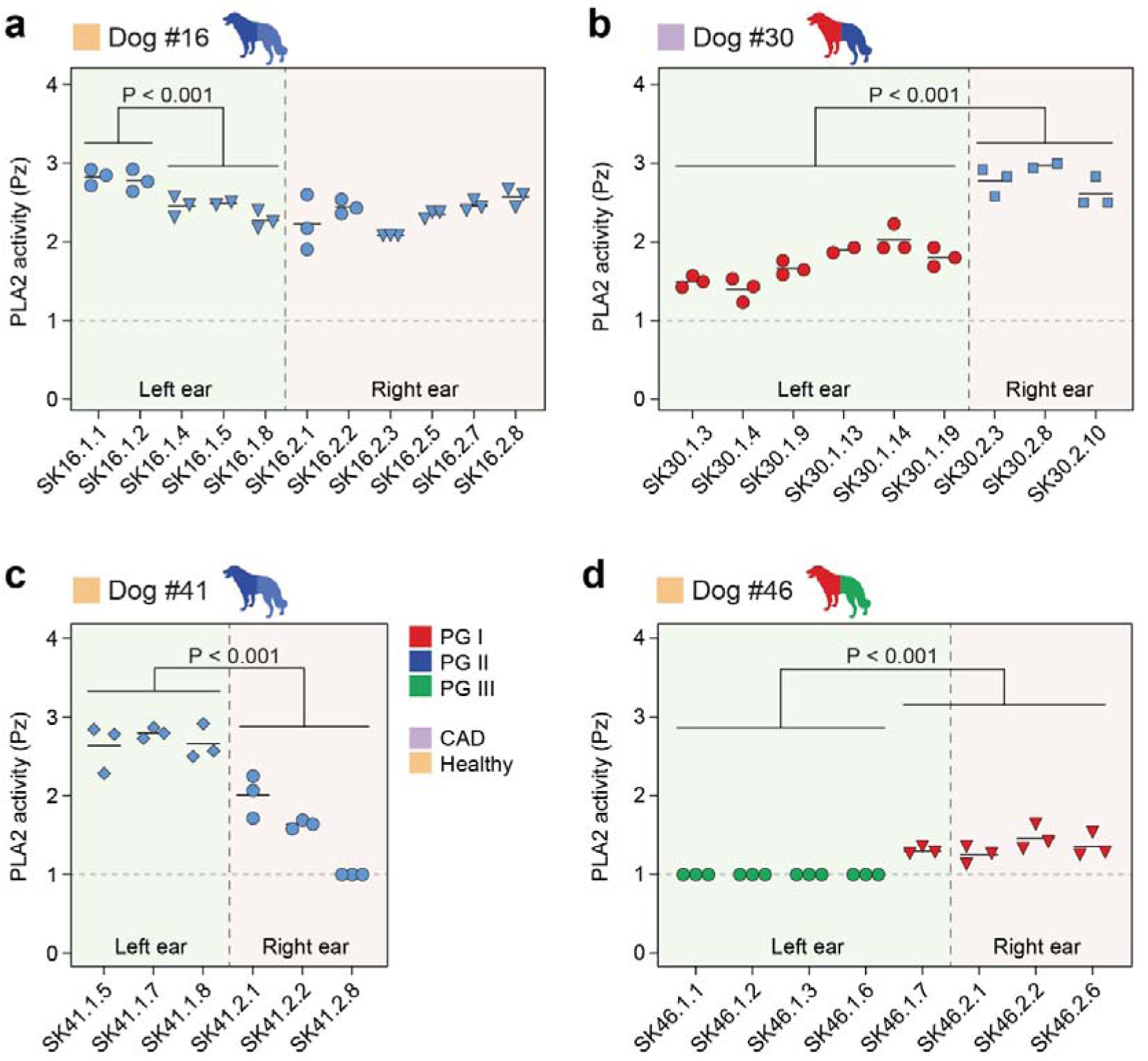
Comparative phospholipase (PLA2) activity among *M. pachydermatis* isolates. PLA2 activity of selected isolates from Dogs #16 (**a**), #30 (**b**), #41 (**c**) and #46 (**d**) was quantified using the Pz value, calculated as the ratio of the total diameter (colony plus precipitation zone) to the colony diameter. Data are the mean Pz values of three replicates per isolate. Statistical differences between groups were determined using unpaired two-tailed t-test with Welch’s correction in panels (**a–c**) and Mann-Whitney test in panel (**d**) (P < 0.05). Symbol color code denotes the three phylogroups, while symbol shapes indicate distinct clusters within each phylogroup (Supplementary Table S8). The vertical dotted line separates isolates recovered from the left and right ears.

### 4.7 Functional correlates of FTIR spectroscopy classification

To assess whether the FTIR spectroscopy-based clustering correlated with any functional traits, we analyzed the subset of isolates subjected to whole-genome sequencing (**Figure 8**, **Supplementary Table S8**) for their phospholipase (PLA2) activity, for which a well-established functional assay is available ^29,30^. Overall, PLA2 activity was generally similar among isolates within the same FTIR cluster or phylogroup but varied significantly between different clusters and phylogroups (**Figure 9**), suggesting a degree of functional differentiation aligned with FTIR spectral profiles. In all tested dogs except Dog #16, isolates from opposite ears exhibited different PLA2 activities. For instance, in Dog #30, right ear isolates (phylogroup II) showed notably higher PLA2 activity compared to those from the left ear (phylogroup I; **Figure 9b**). Intra-ear, intra-cluster, and intra-phylogroup variability were also observed. In Dog #16 (**Figure 9a)**, isolates from the same ear, WGS lineage, and FTIR cluster demonstrated a range of PLA2 activities. Similarly, in Dog #41, isolates from the same phylogroup but different ears also differed in PLA2 activities (**Figure 9c)**, highlighting functional heterogeneity among otherwise closely related strains. A particularly striking example was observed in Dog #46 (**Figure 9d**), where most left ear isolates (phylogroup III) showed low to no PLA2 activity, except for isolate SK46.1.7, which exhibited significantly higher activity. Interestingly, SK46.1.7 belongs to phylogroup I and clustered with right ear isolates, which are also phylogroup I, by both FTIR spectroscopy and WGS. Its enzymatic profile resembled that of the right ear strains, suggesting possible inter-ear transmission and potential involvement in the clinical signs observed in the left ear (**Supplementary Table S1**). Together, these findings suggest that FTIR clustering can capture strain-level functional differences, such as PLA2 activity. However, variability within clusters underscores the importance of integrating FTIR spectroscopy with complementary functional assays to fully resolve phenotypic diversity.

## 5 DISCUSSION

*M. pachydermatis* exhibits substantial intraspecies diversity, which has remained underexplored in the context of CAD. This study applied FTIR spectroscopy to investigate the population dynamics and diversity of *M. pachydermatis* in ears of healthy and CAD-affected dogs and validated the findings using WGS. The observed distribution of distinct phylogroups and differences between healthy and diseased dogs provide new insights into the microbial ecology of *M. pachydermatis* and its potential role in CAD pathogenesis.

*M. pachydermatis* was more easily recovered from ears of CAD-affected compared to healthy dogs, in alignment with earlier published work ^34,35^, and consistent with previous reports of fungal overgrowth in CAD ^36–38^. FTIR spectroscopy-based clustering of the study isolates revealed interesting patterns of heterogeneity and phylogroup distribution. While some dogs had the same lineage and/or phylogroups in both ears, others harbored distinct lineages and/or phylogroups, suggesting either independent colonization events or diversification within the host. Phylogroup II was overall the most common phylogroup in our study population, while phylogroups I and III were disproportionately overrepresented in CAD-affected dogs, indicating potential phenotypic/functional differences between fungal isolates that might influence disease susceptibility and progression. Rather than indicating strict pathogenic specialization, these patterns may reflect differential fitness of specific lineages under inflammatory conditions. Conversely, healthy ears likely maintain a more competitive microbiota environment that limits yeast overgrowth and diversification ^39,40^. The observed variability within individual dogs further highlights the importance of sampling both ears independently and analyzing multiple isolates per ear to capture the full complexity of the colonization profile.

*M. pachydermatis* plays an important role in CAD, where overgrowth of the yeast can amplify skin inflammation through enhanced allergen exposure and modulation of cutaneous immune responses ^16,41^. Thus, assessment of *M. pachydermatis* in CAD should consider not only its presence but also its diversity and the phenotypic traits that may influence pathogenic potential. The trends toward more diverse *M. pachydermatis* populations in CAD dogs, along with the shift in phylogroup abundance in these dogs, suggests an association between disease status and lineage composition, potentially reflecting differential persistence or enrichment of specific lineages in inflamed skin microenvironment. These patterns may arise from repeated recolonization events or within-host evolution, facilitated by skin barrier impairment and/or immune dysregulation in diseased dogs. This hypothesis warrants further investigation into phenotypic traits specific to isolates from CAD dogs. Our findings align with those of Czyzewska et al. ^42^, who analyzed the internal transcribed spacer ITS1 region of nuclear rDNA in *M. pachydermatis* from dogs with and without otitis externa and found that strains from dogs with otitis externa tended to cluster together, though not exclusively. However, the differences in fungal distribution between healthy and CAD dogs were not absolute, emphasizing that besides *M. pachydermatis* diversity other factors also contribute to pathogenesis. A limitation of our study is the fact that although CAD dogs were stratified for *Malassezia* otitis, they were not stratified based on sensitization to *Malassezia*, which may have introduced heterogeneity within the CAD group. Future studies incorporating functional analyses such as metatranscriptomics or metaproteomics may reveal differences in microbial activity that are not detectable through taxonomic profiling alone.

Our results highlight FTIR spectroscopy as an efficient, high-throughput and cost-effective tool for microbial classification in epidemiological and clinical research. For assessing microbial diversity and strain clustering it represents an attractive alternative to WGS. Although it does not have the maximal resolution provided by WGS, FTIR spectroscopy discriminates accurately between closely related isolates within the species of *M. pachydermatis*. Clustering isolates into biologically meaningful groups is largely consistent with WGS-based approaches, highlighting its potential as a frontline technique for screening and resource-efficient prioritization of isolates for downstream sequencing. Importantly, FTIR spectroscopy can also serve as a dereplication method to identify subtle differences among isolates across different samples and/or time points. The ability of FTIR spectroscopy to detect subtle differences in clustering patterns emphasizes its potential for monitoring changes in strain composition and lineage representation over time in relation to clinical outcomes. While this study focused on *M. pachydermatis*, FTIR spectroscopy may be comparably suitable to study the population diversity and lineage-level dynamics in other *Malassezia* spp., including those prevalent in human skin and associated with cutaneous and extracutaneous disorders in humans.

Although the FTIR spectroscopy-based ANN classifier performed well overall, a subset of 85 isolates remained unclassifiable. This could indicate that the model requires further training using a more diverse strain set to better capture the breadth of intraspecific and lineage-level variability. Expanding the training dataset may improve the classifier’s generalizability and its ability to detect currently underrepresented or atypical strains. Additionally, the phylogroup classifier was developed using haploid strains, and it is possible that isolates with ambiguous phylogroup assignments are diploid or aneuploid, which could affect the classifier’s performance. The presence of hybrid isolates also cannot be excluded and may reflect genetic exchange facilitated by the retention of mating-type loci ^28^, although sexual reproduction has not yet been directly observed in *Malassezia* species. Such genetic complexity could contribute to intermediate or ambiguous FTIR profiles, highlighting the need to consider ploidy variation and potential hybridization events in future FTIR-classification models.

This study also highlights the importance of selecting appropriate sampling techniques and sites to accurately capture the microbial diversity in a relevant clinical context, given the observed niche-specific differences in *M. pachydermatis* assemblages. While superficial swabs may identify microbial communities not sampled from deeper locations, deep ear swabs provide a more representative snapshot of the disease-relevant compartment.

Together, this study provides a foundation for understanding the population structured diversity of *M. pachydermatis* in both health and disease scenarios and offers new insights into the microbial dynamics associated with CAD. By focusing on the FTIR spectral region that corresponds to polysaccharides, the detected strain-level variations may reflect subtle differences in the yeast’s carbohydrate profile, with potential relevance for the crosstalk of *M. pachydermatis* with the host, given the prominent role of fungal carbohydrates as host recognition targets ^43,44^. The observed shifts in fungal lineage composition and diversity underscore the complex interplay between host factors and microbial adaptation in atopic environments. Future research should focus on uncovering the genetic and metabolic determinants underlying homeostatic vs. pathogenic fungus-host interactions and on clarifying how these processes shape lineage persistence and turnover within atopic environments. As such, integrating FTIR spectroscopy with other omics approaches, such as genomics, transcriptomics or metabolomics, has the potential to provide a more comprehensive understanding of *M. pachydermatis* biology and its role in CAD. Furthermore, the development of FTIR spectroscopy-based predictive models for identifying pathogenic phenotypes could significantly enhance its application in both clinical and research settings. Ultimately, these findings contribute to a deeper understanding of the role of microbial diversity in health and disease, with broad implications for both veterinary and human medicine.

## 6 ACKNOWLEDGMENTS

The authors would like to thank Roger Stephan and members of the LeibundGut, Favrot and Heitman groups for helpful advice and discussion. This work was supported by the Vetsuisse Faculty of University of Zurich, the Swiss National Science Foundation (grant # 320030-227692 to SLL), the National Institute of Health (NIH/NIAID R21 AI168672-02 to SLL and JH). JH is Co-Director and Fellow of the CIFAR program Fungal Kingdom: Threats & Opportunities.

## 7 AUTHOR CONTRIBUTIONS

**SK**: conceptualization, methodology, formal analysis, investigation, writing – original draft

**MAC**: formal analysis, data curation, writing – review & editing

**SM**: investigation, writing – review & editing

**AR**: conceptualization, supervision, writing – review & editing, funding acquisition

**NF**: supervision, writing – review & editing, funding acquisition

**FM**: supervision, writing – review & editing,

**MK**: investigation, writing – review & editing

**MDP**: formal analysis, data curation, writing – review & editing

**JH**: resources, writing – review & editing

**SLL**: conceptualization, supervision, writing – review & editing, visualization, project administration, funding acquisition

**CF**: conceptualization, supervision, writing – review & editing, project administration, funding acquisition

**FM**: conceptualization, methodology, validation, formal analysis, investigation, writing - original draft, visualization,

## 8 COMPETING INTERESTS

The authors have no competing interests to declare.

## Notes

### Competing Interest Statement

The authors have declared no competing interest.

