## Supplementary material for "Fourier transform infrared spectroscopy reveals high intraspecies diversity of *Malassezia pachydermatis* in dogs with atopic dermatitis": Supllementary Figures

**Supplementary Figures**

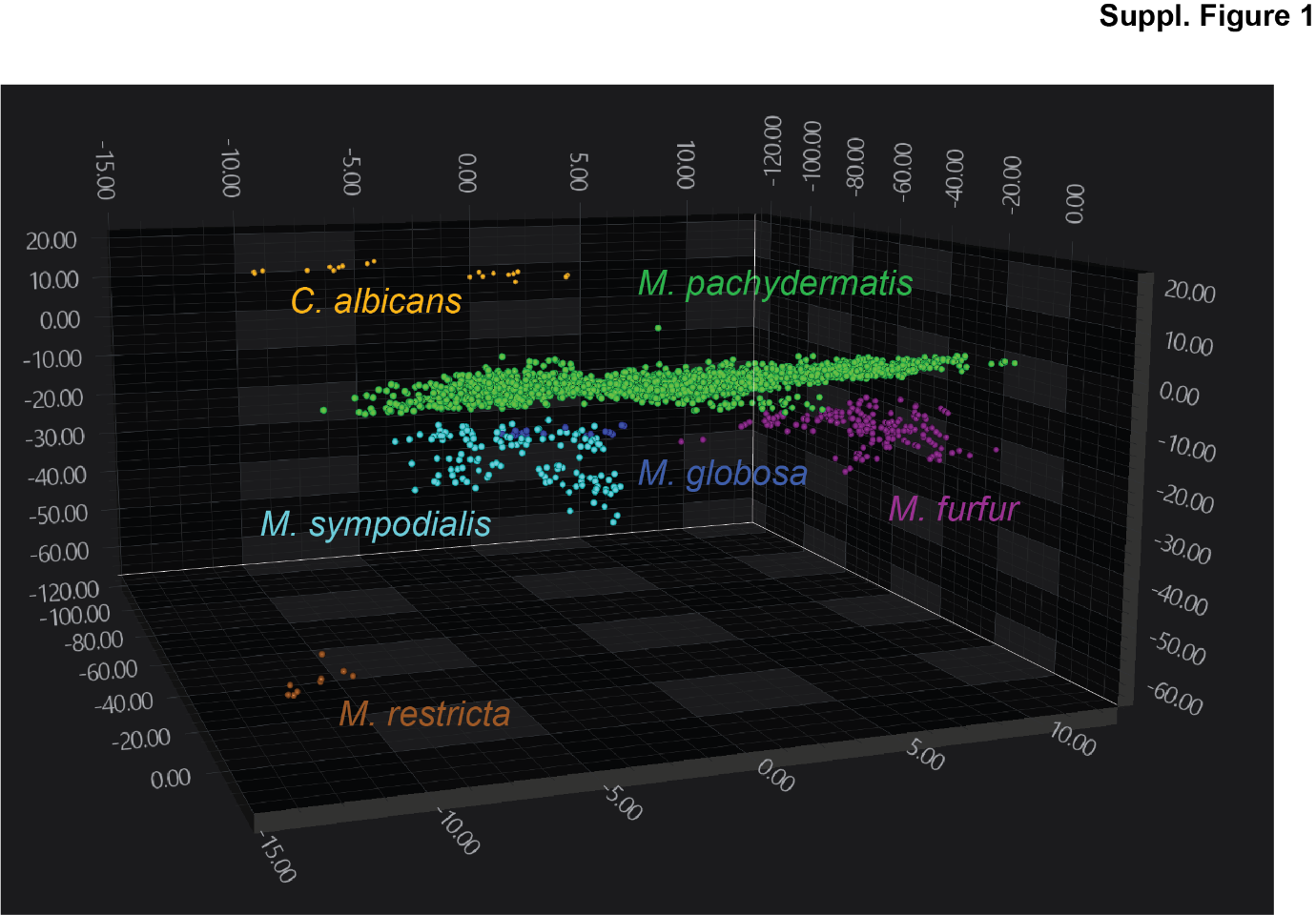

**Supplementary Figure S1. Separation of *M. pachydermatis* from other *Malassezia* species in the FTIR spectral space.** As Figure 2, but view from a different angle. 3D scatter plot showing the distribution of *M. pachydermatis* isolates from study dog ears (n = 539) and reference strains of *Malassezia furfur* (n = 12), *M. globosa* (n = 2), *M. sympodialis* (n = 6), *M. restricta* (n = 1) and *C. albicans* (n = 2) within the FTIR spectral space. The plot displays the first three LD axes. Each isolate is represented by at least six spectra, with each symbol (•) corresponding to a technical replicate from at least two independent biological repeats, i.e., each isolate is represented by at least 6 symbols. Dimensionality reduction was performed using LDA.

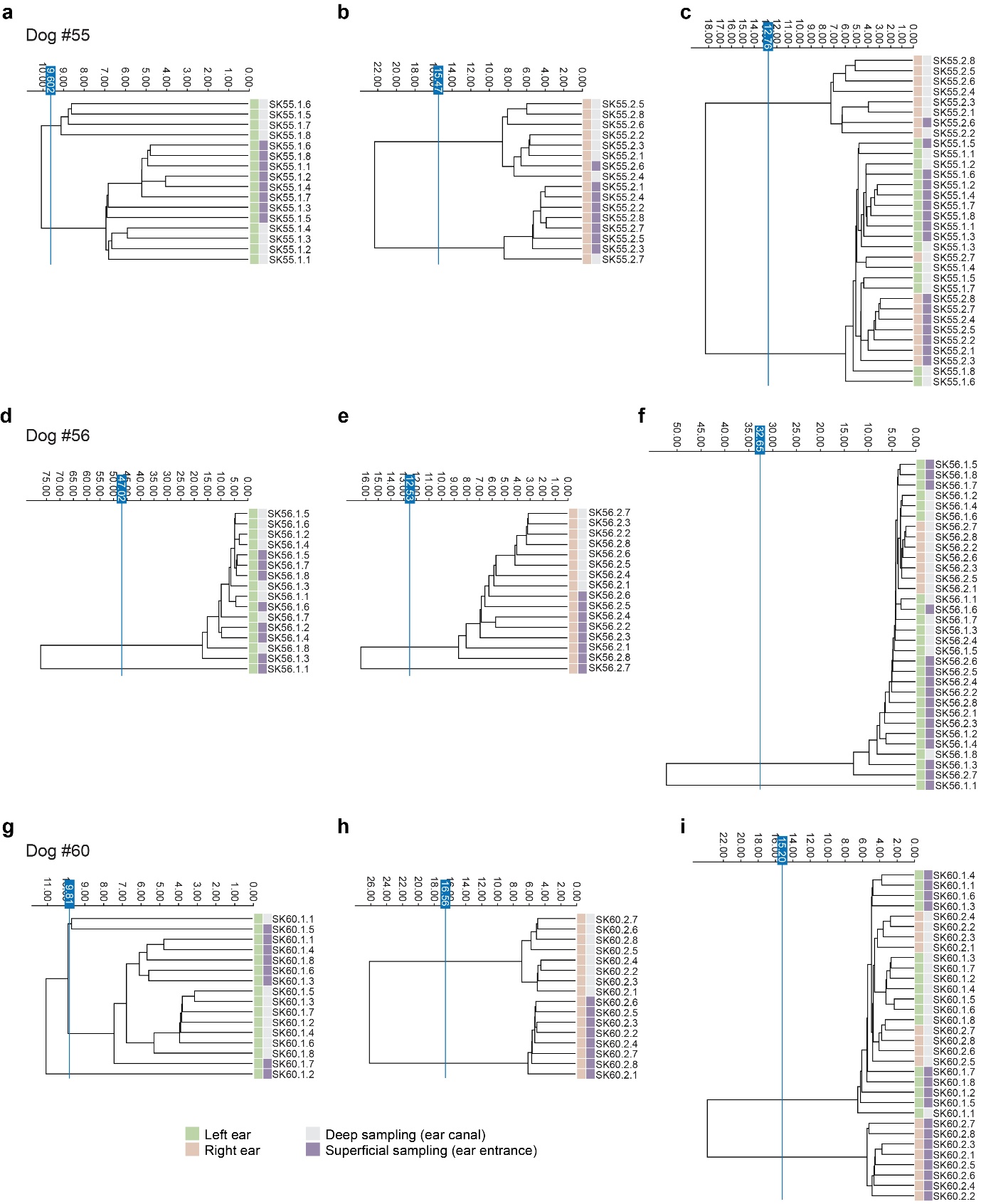

**Supplementary Figure S2**. **Clustering of *M. pachydermatis* isolates from Dogs #55, #56, and #60 in the FTIR spectral space.** Dendrograms generated from clustering FTIR spectra of *M. pachydermatis* isolates collected from (**a, d, g**) the left ear, (**b, e, h**) the right ear, and (**c, f, i**) both ears of Dogs #55 (**a**–**c**), #56 (**d**–**f**), and #60 (**g**–**i**). Samples were collected twice on the same day either by superficial swabbing (purple) or deep ear swabbing (light grey). Each row represents the average spectrum derived from at least two independent biological replicates. The vertical blue line indicates the automatically calculated clustering cut-off. Dimensionality reduction was performed using LDA. The analysis focused on the wavenumber region 1300–800 cm⁻¹, corresponding to polysaccharides.

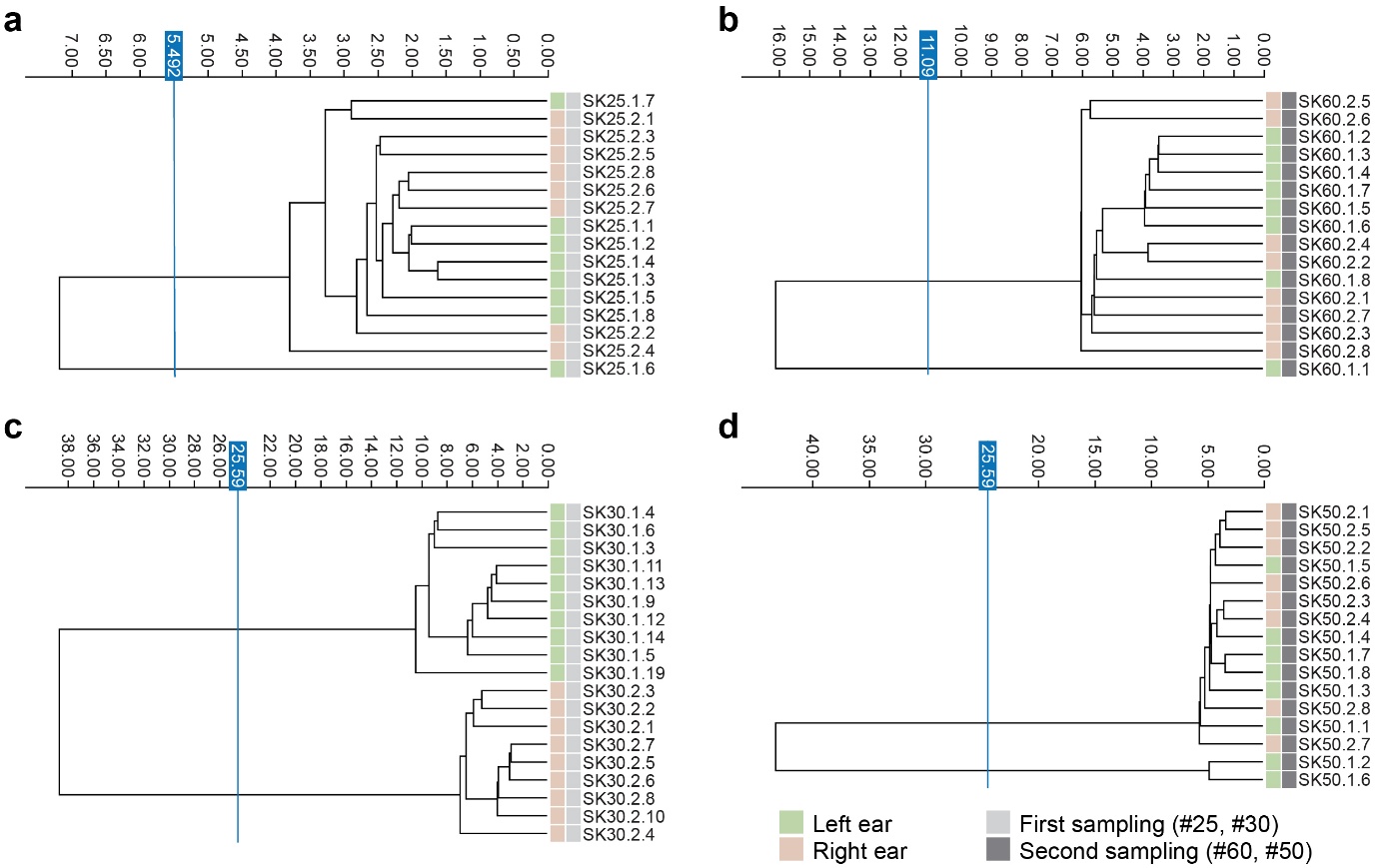

**Supplementary Figure S3. Shifts in *M. pachydermatis* populations over time.** Dendrograms generated from clustering FTIR spectra of *M. pachydermatis* isolates derived from a healthy dog (**a–b**; Dog #25) and a CAD-affected dog (**c–d**; Dog #30), each sampled twice over an interval of more than two months. (**a**) First sampling of Dog #25 (#25), (**b**) second sampling of Dog #25 (#60), (**c**) first sampling of Dog #30 (#30), (**d**) second sampling of Dog #30 (#50). Clustering was based on the average spectra from at least two independent biological replicates, each with three technical replicates. The vertical blue line denotes the automatically calculated clustering cut-off. Dimensionality reduction was performed using LDA. The analysis focused on the wavenumber region 1300–800 cm⁻¹, corresponding to polysaccharides.

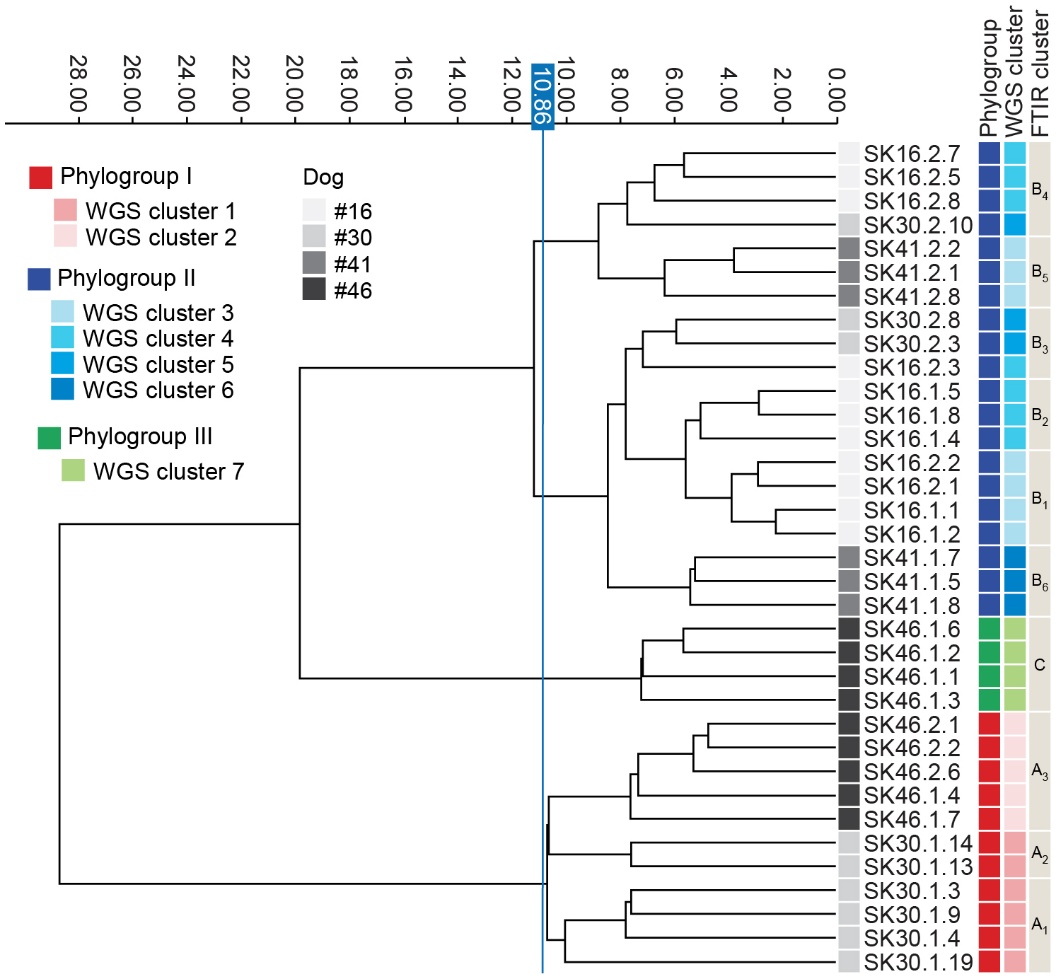

**Supplementary Figure S4. Clustering of sequenced *M. pachydermatis* isolates from study dogs in the FTIR spectral space.** Dendrograms generated from clustering FTIR spectra of sequenced *M. pachydermatis* isolates collected from Dog #16, Dog #30, Dog #41, and Dog #46. Each row represents the average spectrum derived from at least two independent biological replicates. The vertical blue line indicates the clustering cut-off. Dimensionality reduction was performed using LDA. Spectral analysis focused on the wavenumber region 1300–800 cm⁻¹ region, corresponding to polysaccharides. The WGS based phylogroups and clusters are also indicated.
